# Adipose tissue dysfunction and visceral fat are associated to hepatic insulin resistance and severity of NASH even in lean individuals

**DOI:** 10.1101/2022.01.19.476711

**Authors:** Chiara Saponaro, Silvia Sabatini, Melania Gaggini, Fabrizia Carli, Chiara Rosso, Vincenzo Positano, Angelo Armandi, Gian Paolo Caviglia, Riccardo Faletti, Elisabetta Bugianesi, Amalia Gastaldelli

**Affiliations:** Cardiometabolic Risk Unit, Institute of Clinical Physiology, CNR, Pisa, Italy; Division of Gastroenterology and Hepatology and Lab. of Diabetology, Dept. of Medical Sciences, University of Turin, Turin, Italy; Fondazione Toscana Gabriele Monasterio, Pisa, Italy

**Author notes:** **Correspondence to:** Amalia Gastaldelli, PhD, Head of Cardiometabolic Risk Unit, Institute of Clinical Physiology, CNR, via Moruzzi 1 56100 Pisa Italy, /80, Chiara Saponaro, PhD, University of Lille, CHU Lille, Inserm U1190, EGID, Faculté de Médecine de Lille, Pôle Recherche, 1, Place de Verdun, 59045 - Lille, France. Univ. Lille, CHU Lille, Inserm U1190, EGID, Lille, France.

**Keywords:** NAFLD, fat distribution, visceral fat, subcutaneous fat, insulin resistance

## Abstract

**Background & Aims:** Nonalcoholic fatty liver disease (NAFLD) is a heterogeneous disorder, but the factors that determine this heterogeneity remain poorly understood. Adipose tissue (AT) dysfunction is causally linked to NAFLD since it causes intrahepatic triglyceride (IHTG) accumulation through increased hepatic lipid flow, due to insulin resistance (IR) and pro-inflammatory adipokines release. While many studies in NAFLD have looked at total adiposity (that is mainly subcutaneous fat, SC-AT), it is still unclear the impact of visceral fat (VF). Thus, we investigated how VF vs. SC-AT were related to NAFLD in lean, overweight, and obese individuals compared to lean controls.

**Methods:** Thirty-four non-diabetic NAFLD with liver biopsy and eight lean control individuals (CT) were enrolled in this study. We measured fat distribution (VF, SC-AT and IHTG) by magnetic resonance imaging (MRI), adiponectin concentration, free fatty acids (FFAs) and triglyceride (TAG) concentration and composition by mass spectrometry (MS), lipolysis and IR by tracer infusion.

**Results:** IHTG was positively associated with lipolysis, adipose tissue IR, TG concentrations, and increased ratio of saturated/unsaturated fatty acids. VF was higher in NAFLD (including lean individuals) compared to controls, was increased with fibrosis stage and was associated with IR in liver, muscle and adipose tissue, increased lipolysis, and decreased adiponectin levels. Collectively, our results suggest that VF accumulation, given its location close to the liver, is one of the major risk factors for NAFLD.

**Conclusions:** These findings propose VF as an early indicator of NAFLD independently of BMI, which may allow for evidence-based prevention and intervention strategies.

## Introduction

Non-alcoholic fatty liver disease (NAFLD) is a highly heterogeneous metabolic disorder covering a wide spectrum of liver diseases that ranges from simple steatosis to progressive inflammation, fibrosis, and ballooning, resulting in non-alcoholic steatohepatitis (NASH) and cirrhosis [1-3]. The pathophysiology of the onset and progression of NAFLD is complex and is still not completely understood although it is established that NAFLD is a metabolic disease. In the last decades, a “multiple parallel hits hypothesis” has been proposed, in which not only excess calories and dietary components such as the westernized diet favor hepatic triglyceride accumulation and de novo fatty acid synthesis, but also the interactions between the liver and adipose tissue, together with the gut, play a concomitant role in the development of the disease. Although the VF depot is much smaller than the SC-AT it provides a foreground effect, by driving the release of FFAs directly into the portal circulation [2, 4, 5], possibly due to its enhanced lipolytic activity compared to that of SC-AT [6]. Indeed, the accumulation of lipids in the VF is often complemented with storage in other ectopic sites, such as the liver and the pancreas [7, 8]. Moreover, hepatic and circulating lipid composition may also affect the extent of liver damage, inflammation, and the rate of disease progression [9-12]. Finally, in this complex scenario, hepatic and peripheral (muscle and adipose tissue) insulin resistance are central, although a causal link between steatosis, VF accumulation and insulin resistance requires more investigation. Although various *in vivo*, ex-situ and *in vitro* models have been extensively used to study the role of VF and insulin resistance in NAFLD [13-16], these models don’t represent the full spectrum of human NAFLD progression [17]. By contrast, numerous human studies, despite their translational value, are highly heterogeneous, due to ethnicity, sex, age, and the presence of confounding factors such as concomitant metabolic diseases, with and without drug treatment. Moreover, these studies are often confronted with limiting factors in the methodology such as the accurate measurement of parameters including fat distribution and insulin resistance. Therefore, the use of stable-isotope methodology provides a more thorough mechanistic readout of lipid metabolism and insulin resistance [18, 19]. Collectively, these techniques combined with the diagnosis and staging of NAFLD by non-invasive imaging techniques [20, 21], will unveil whether or not VF accumulation and dysfunction are the initiators of NAFLD onset and progression. Upon recruitment of a cohort of non-diabetic Italian patients with various stages of NAFLD, stratified for BMI, we set out to determine the interplay of VF and insulin resistance at different stages of NAFLD. Indeed, we forecasted that this subset of patients would offer a unique setting to assess the direct impact of fat distribution and dysfunction on hepatic insulin resistance and steatosis, independently of diabetes, an area that has not been adequately investigated.

## Materials and Methods

### Patients

Thirty-four non-diabetic subjects, with biopsy-proven NAFLD and eight controls (CT) were enrolled for the study that was part of the activities of the EU-FLIP project. Liver biopsies were scored according to Kleiner et al [22], altered fat topography, liver and pancreatic fat amount were measured by magnetic resonance imaging (MRI). The study was approved by the ethics committee of the University Hospital San Giovanni Battista of Torino and was in accordance with the Helsinki Declaration. All subjects signed an informed consent form before participating to the study.

### Plasma measurements

In all subjects we measured fasting plasma lipid profile, concentrations of FFA, triglycerides, total and HDL cholesterol, LDL, LFTs, apolipoprotein B (APO-B), apolipoprotein A1 (APO-A1), glucose, insulin and C-peptide, as previously reported [23, 24]. We also measured circulating concentrations of markers of inflammation and oxidative stress, e.g., MCP-1, and circulating adiponectin by Multiplex Assay based on Luminex Technologies.

### Liver damage and fibrosis score

Liver biopsy was scored according to Kleiner et al [22]. The sum of grading for steatosis, lobular inflammation and hepatocellular ballooning was used to calculate the NAFLD activity score (NAS) from 0 to 6. Fibrosis was staged 0 to 4 and classified as absent (0), mild (1-2) and severe (3-4). NASH was defined by the local pathologist according to the joint presence of steatosis, hepatocyte ballooning and lobular inflammation with or without fibrosis.

### Fat topography by magnetic resonance imaging

Magnetic resonance images were acquired using the DIXON method following protocols previously published [25, 26]. MRI images were acquired in DICOM format and then transferred to a dedicated workstation to be analyzed by an ad hoc developed software [27]. For each slice the subcutaneous abdominal fat area was measured by automatic detection of the outer and inner margins of subcutaneous adipose tissue as region of interest (ROI) from the cross-sectional images, and by counting the number of pixels between the outer and inner margins of subcutaneous adipose tissue; volume was automatically calculated by multiplying the slice area by the slice thickness and then the values of all slices were summed to obtain the abdominal fat volume. For each slice visceral (intra-abdominal) fat area and volume was determined with the use of histograms specific to the visceral region as previously described [27] and then summed to obtain the abdominal VF volume. A factor of 0.92 was used to convert adipose tissue volume into adipose tissue mass [25]. To quantify liver fat fraction (LFF) the reconstructed fat and water image were analyzed using the OsiriX program [28]; 3 regions of interest (ROI) (30 × 30 mm) were quantified in the reconstructed fat and water images. Fat content was calculated from signal intensity in the ROIs of the reconstructed fat (S_fat_) and water (S_water_) images as [S_fat_/(S_water_+S_fat_) × 100] [26]. Fat content was considered normal in liver if < 5.6% [29].

### Metabolic indexes

Endogenous glucose production (EGP) and peripheral glucose clearance were measured by the kinetics of 6,6-D2-glucose infused for 2 hours during a fasting state, following a protocol previously reported [23, 30]. Hepatic insulin resistance (Hep-IR) reflects the inability of insulin to suppress fasting EGP and was calculated as the product of EGP x fasting insulin [24] while Muscle-IR was measured as the inverse of peripheral glucose clearance normalized by fasting insulin [30]. Lipolysis was measured by the kinetics of D5-glycerol infused for 2 hours during a fasting state, following a protocol previously reported [23, 30]. Adipose tissue insulin resistance reflects the inability of insulin to exert an antilipolytic action, leading to a relative increase in lipolysis and fasting FFA concentrations [31]. Lipo-IR was calculated as the product of fasting lipolysis x insulin while Adipo-IR was calculated as the product of fasting FFA x insulin [23, 24, 31].

### Metabolite analysis

Free fatty acid composition was evaluated by GC-MS (Agilent technology GC7890-MS5975). Briefly, separation of lipid fraction from 20uL of plasma was carried out with a methanol and chloroform (2:1) solution using the Folch’s method [32]. Fatty acids were derivatized to methyl-ester with a solution of methanol-BF3 14% and dried. Samples were reconstituted with 70uL heptane and 1uL was injected in the GC-MS. Total plasma FFA concentration was measured spectrophotometric ally on Beckman analyzer (Waco, Global Medical Instrumentation, Ramsey, MN). Single fatty acid concentration was determined using as internal standard a mix of 13C labeled FFA (Alga mix, CIL Cambridge MA, USA) and heptadecanoic acid. We quantified saturated (SFA, myristic, palmitic and stearic acid), and unsaturated (UFA, i.e., the mono-unsaturated oleic and palmitoleic acids and the poly-unsaturated linoleic and arachidonic acids) fatty acid concentrations. The ratio of SFA/UFA were calculated as parameters of lipotoxicity. Triglycerides (TAGs) plasma composition was analyzed by ultra-high-pressure liquid chromatography/quadrupole time-of-flight mass spectrometry (UHPLC/Q-TOF, 1290 Infinity-6540 Agilent Technology, Santa Clara, CA) equipped with electrospray ionization (ESI). Plasma (20µL) was deproteinized with 200µL of cold methanol (Merck, Darmstadt, Germany), centrifuged at 14000 rpm for 20 minutes and the supernatant was transferred into glass vials and 1 µL was injected into the LC/MS. Triglycerides were separated used an Agilent ZORBAX Eclipse Plus C18 2.1 × 100 mm 1.8-Micron and acquisition was set in positive mode. TG species were analyzed using Agilent MassHunter Profinder B.06.00. Analysis of TG composition was qualitative: area of each specie was normalized to total area of triglycerides. TGs were classified as low unsaturated by calculating sum of areas of TGs with 0 and 1 double bonds (0-1 db) and unsaturated by calculating sum of areas of TGs with 2-5 double bonds (2-5 db).

### Statistical analysis

Variables were expressed as mean +/- standard error (SE). Differences in distribution among groups were calculated using non parametric Mann-Whitney’s test. Correlation analysis was performed using Pearson’s correlation coefficient, after a log transformation of variables.

Heatmaps were created reporting data as median within the groups of interest. Data were therefore centered and scaled in the row direction to improve interpretability. All statistical analysis was performed using R Statistical Software (version 4.0.5).

## Results

### 1. Clinical characteristics of study subjects

We studied thirty-four subjects with biopsy-proven NAFLD (20 NASH and 12 NAFL) with a wide range of BMI (n=10 with BMI <25; n=12 with BMI ≥ 25 and <30; n=12 with BMI ≥30) and eight lean controls (**Fig. 1A**). Fasting glycemia, circulating FFAs and high-density lipoprotein (HDL) were not significantly different among the 4 groups (**Table 1**). Plasma levels of liver enzymes were significantly higher in NAFLD compared to the control group, i.e. alanine aminotransferase (ALT: 75.0±6.4 vs 16.3±1.6 U/l), aspartate aminotransferase (AST: 45.7±6.2 vs 19.6±1.6 U/l) and gamma-glutamyltransferase (GGT: 105.9±21.2 vs 14.3±4.2 U/l, all p<0.0002). Total TG, low-density lipoprotein (LDL) and cholesterol were higher in NAFLD subjects compared to the controls.

**Table 1:**
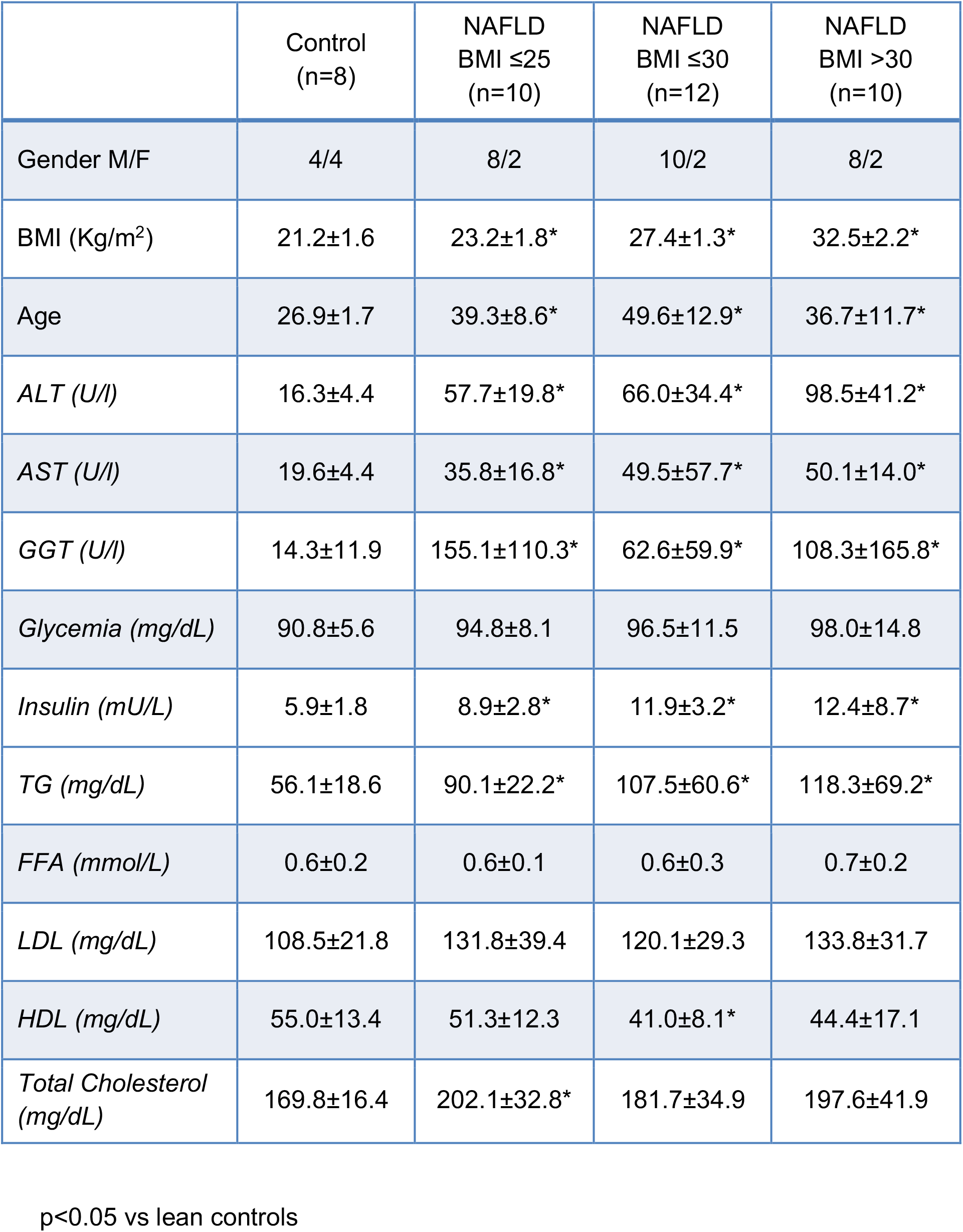
Clinical Characteristics of study subjects.

|  | Control<br>(n=8) | NAFLD<br>BMI ≤25<br>(n=10) | NAFLD<br>BMI ≤30<br>(n=12) | NAFLD<br>BMI >30<br>(n=10) |
| --- | --- | --- | --- | --- |
| Gender M/F | 4/4 | 8/2 | 10/2 | 8/2 |
| BMI (Kg/m<sup>2</sup>) | 21.2±1.6 | 23.2±1.8* | 27.4±1.3* | 32.5±2.2* |
| Age | 26.9±1.7 | 39.3±8.6* | 49.6±12.9* | 36.7±11.7* |
| *ALT (U/l)* | 16.3±4.4 | 57.7±19.8* | 66.0±34.4* | 98.5±41.2* |
| *AST (U/l)* | 19.6±4.4 | 35.8±16.8* | 49.5±57.7* | 50.1±14.0* |
| *GGT (U/l)* | 14.3±11.9 | 155.1±110.3* | 62.6±59.9* | 108.3±165.8* |
| *Glycemia (mg/dL)* | 90.8±5.6 | 94.8±8.1 | 96.5±11.5 | 98.0±14.8 |
| *Insulin (mU/L)* | 5.9±1.8 | 8.9±2.8* | 11.9±3.2* | 12.4±8.7* |
| *TG (mg/dL)* | 56.1±18.6 | 90.1±22.2* | 107.5±60.6* | 118.3±69.2* |
| *FFA (mmol/L)* | 0.6±0.2 | 0.6±0.1 | 0.6±0.3 | 0.7±0.2 |
| *LDL (mg/dL)* | 108.5±21.8 | 131.8±39.4 | 120.1±29.3 | 133.8±31.7 |
| *HDL (mg/dL)* | 55.0±13.4 | 51.3±12.3 | 41.0±8.1* | 44.4±17.1 |
| *Total Cholesterol (mg/dL)* | 169.8±16.4 | 202.1±32.8* | 181.7±34.9 | 197.6±41.9 |

p<0.05 vs lean controls

**Figure 1.**
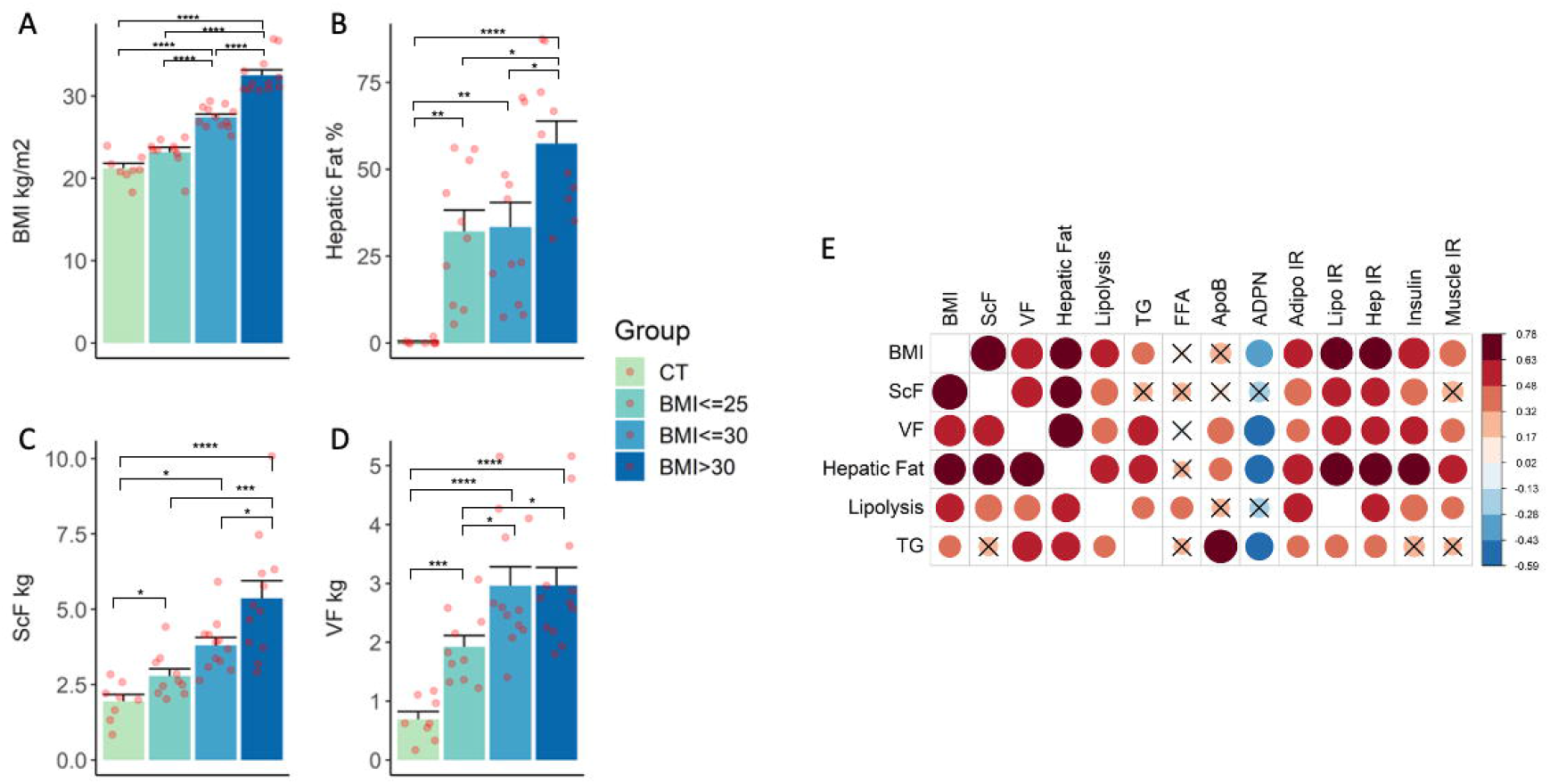
BMI (panel A) and intra hepatic triglyceride content measured by MRI, i.e., IHTG (panel B), subcutaneous fat (panel C), and visceral fat (panel D), in healthy controls and NAFLD subjects divided in accordance with their BMI (from light green to dark blue). Data are expressed as means ± SE, differences among the groups were tested using non-parametric Mann Whitney’s test. Specific p values: *<0.05, ** <0.01, ***<0.001, ****<0.0001. Panel E shows the univariate correlation matrix between BMI, fat distribution, lipolysis and circulating TG and metabolic parameters in the entire cohort. Correlations that are not statistically significant (Pearson’s p-value≥0.05) were crossed out.

### 2. Obesity and adipose tissue distribution in NAFLD

Subjects with NAFLD were divided into 3 groups according to their BMI as lean (BMI <25 kg/m^2^), overweight (BMI ≥ 25 and <30 kg/m^2^) and obese (BMI ≥ 30 kg/m^2^). BMI of lean individuals with NAFLD was similar to healthy lean subjects without NAFLD (23.2±0.6 vs 21.2±0.6 kg/m^2^) (**Fig. 1A**). Excessive intrahepatic triglyceride content (IHTG) (**Fig. 1B**) and visceral fat (VF) (**Fig. 1D**) measured by MRI were increased in all NAFLD compared to the lean controls. SC-AT, on the other hand, was significantly increased only in overweight and obese NAFLD, compared to the lean NAFLD and lean controls (**Fig. 1C**). Obesity (BMI≥30 kg/m^2^) was associated to a further increase in all fat depots (**Fig. 1B-D**).

We analyzed body fat distribution according to the grade of steatosis, lobular inflammation and hepatocellular ballooning (NAS score, two groups NAS≤3 and NAS≥4) (**Fig. 2 A-D**). As expected, hepatic fat increased proportionally with the grade of NAS score, while BMI, VF and SC-AT showed a trend to be higher in NAFLD patients with high NAS score without reaching statistical significance (**Fig. 2 A-D**). In the analysis of body fat distribution according to the degree of fibrosis (**Fig. 3 A-D**) we observed that subjects with higher fibrosis (i.e. F2-F4), had higher BMI, SC-AT and VF compared to F01 and to lean controls (**Fig. 3 A,C,D**), in line with previous studies [33].

**Figure 2.**
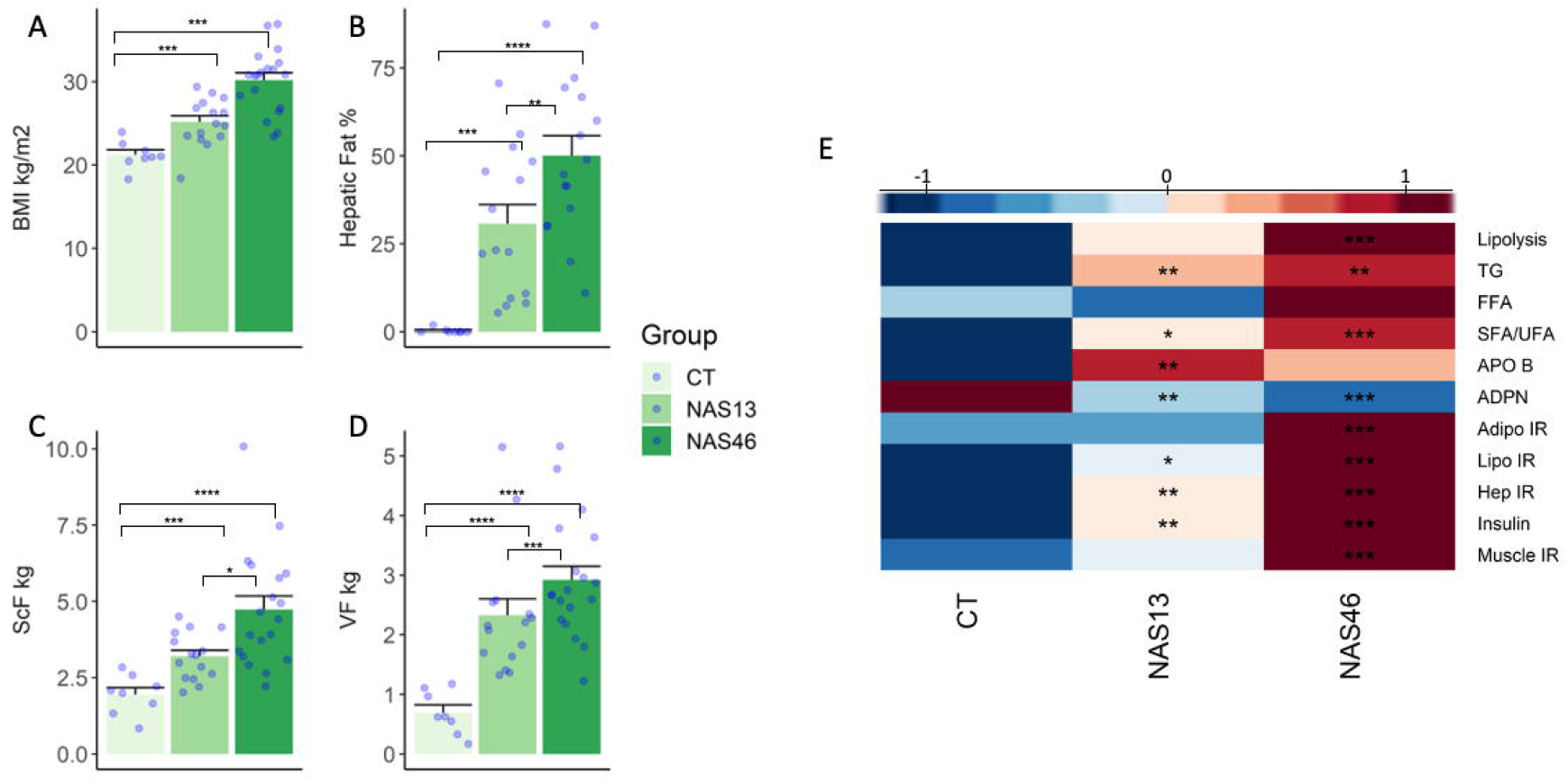
BMI (panel A) and fat distribution measured by MRI, i.e., IHTG (panel B), subcutaneous fat (panel C), visceral fat (panel D), in healthy controls and NAFLD subjects divided in accordance with their grade of steatosis, lobular inflammation and hepatocellular ballooning (NAS score) (from light green to dark green). Data are expressed as means ± SE, non-parametric Mann Whitney’s test was used to test differences among the groups. Specific p values: *<0.05, ** <0.01, ***<0.001, ****<0.0001. In panel E heatmap of metabolic parameters were reported in controls and NAFLD subjects divided by their grade of NAS score. Values are reported as median within the groups and normalization was applied to the rows. Specific p values for non-parametric Mann Whitney’s test vs control subjects: *<0.05, ** <0.01, ***<0.001, ****<0.0001.

**Figure 3.**
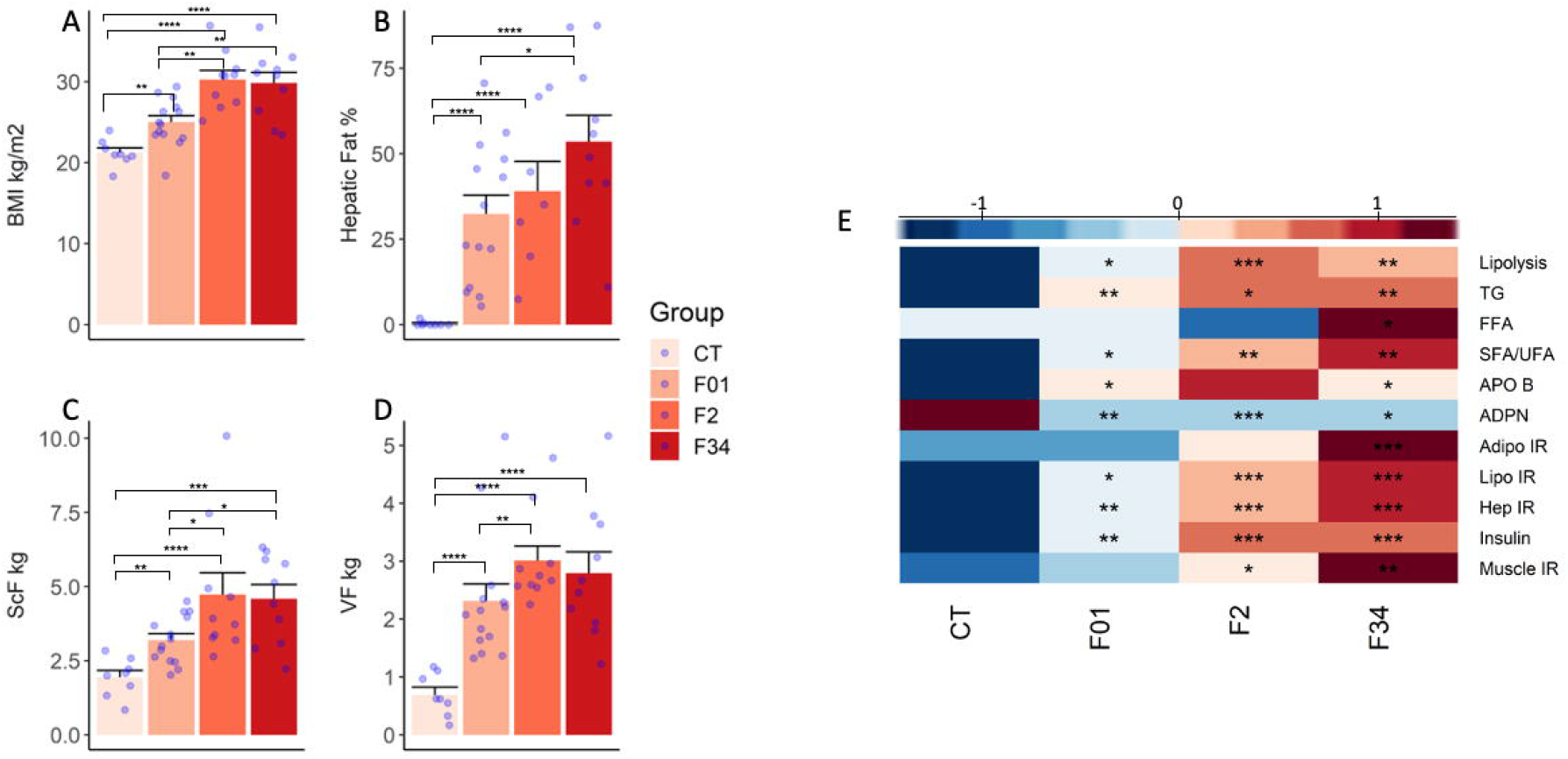
BMI (panel A) and fat distribution measured by MRI, i.e., IHTG (panel B), subcutaneous fat (panel C), visceral fat (panel D), in healthy controls and NAFLD subjects divided in accordance with their degree of fibrosis (from pink to dark red). Data are expressed as means ± SE, non-parametric Mann Whitney’s test was used. Specific p values: *<0.05, ** <0.01, ***<0.001, ****<0.0001. In panel E heatmap of metabolic parameters were reported in controls and NAFLD subjects divided by their degree of fibrosis. Values are reported as median within the groups and normalization was applied to the rows. Specific p values for non-parametric Mann Whitney’s test vs control subjects: *<0.05, ** <0.01, ***<0.001, ****<0.0001.

### 3. Insulin resistance and adipose tissue distribution in NAFLD

Tracer infusion allowed the measurement of metabolic fluxes such as adipose tissue lipolysis and endogenous glucose production (mainly hepatic), and to evaluate insulin resistance in the liver (Hep-IR), muscle (Muscle-IR) and adipose tissue (as Adipo-IR and Lipo-IR). Adipose tissue lipolysis was increased in NAFLD compared to healthy controls and associated not only with BMI, but also with increased abdominal fat VF, SC-AT and IHTG (**Fig. 1E**). Moreover, lipolysis was positively correlated with increased FFA concentrations demonstrating the increased overflow of fatty acids from the periphery to other organs in subjects with NAFLD (**Fig. 1E**). The associations were similar in NAFL and NASH (data not shown), but lipolysis was higher in subjects with more severe forms of NAFLD, i.e., in subjects with NAS ≥4 (**Fig. 2E**) and with fibrosis ≥2 (**Fig. 3E**). Adipose tissue IR that measures the defective action of insulin in suppressing peripheral lipolysis (here measured as Adipo-IR and Lipo-IR) was positively associated with IHTG (**Fig. 1E**) and was higher in NAFLD subjects with a more severe liver disease (**Fig. 2E and 3E**). Of note, VF was also inversely associated with circulating adiponectin levels (**Fig. 1E**), the most abundant anti-inflammatory and atheroprotective adipokine secreted by adipose tissues [34] that is also a marker of insulin sensitivity. However, we did not observe a correlation between SC-AT and adiponectin levels in line with previous studies [35, 36]. It has been already established that adiponectin reduces hepatic gluconeogenesis and increases glucose uptake and whole-body insulin sensitivity [37, 38]. Here, we found that adiponectin was significantly decreased in all NAFLD individuals compared to controls also in those with low BMI (p < 0.02), and it was inversely correlated with Hep-IR (rho= -0.50, p<0.001), and IHTG accumulation (**Fig. 1E**), thus posing the possibility of an additional mechanism by which VF negatively impacts liver function and promotes NAFLD. Endogenous glucose production was slightly higher in subjects NAFLD compared to lean controls (14.2±0.3 vs 11.7±0.6 µmol/min per Kg of fat free mass, p=0.001) including in lean NAFLD (p=0.02). Hep-IR and Muscle-IR were increased in NAFLD subjects, especially in those with a more severe liver disease, i.e., in subjects with NAS ≥4 (**Fig. 2E)** and with fibrosis ≥2 (**Fig. 3E**). VF and SC-AT were strongly correlated with Hep-IR, and with a less extent to Muscle-IR (**Fig. 1E**), as previously shown [2].

### 4. Lipid concentrations and composition, and adipose tissue distribution in NAFLD

Although BMI was associated to increased lipid concentrations, only VF and IHTG, not SC-AT, were associated to increased circulating TG and ApoB concentrations (**Fig. 1E**) indicating increased synthesis and secretion of VLDL, likely stimulated by a high portal lipid supply given that VF is more lipolytic than SC-AT [6, 39, 40].

The analysis of FFA and TG composition showed a shift towards increased saturated FFAs and TGs in NAFLD vs CT, although the increase was more pronounced in subjects with BMI≥30 (**Fig. 4 A,C**). The severity of liver disease (i.e., higher fibrosis) was associated to higher circulating saturated fatty acids both as FFA (**Fig. 4 B**), such as myristic, palmitic and stearic acid, with a depletion of linoleic acid, as well as higher saturated TG (**Fig. 4 D**) in line with previous studies that shown that subjects with NAFLD have higher intrahepatic and circulating lipids containing saturated fatty acids [41].

**Figure 4.**
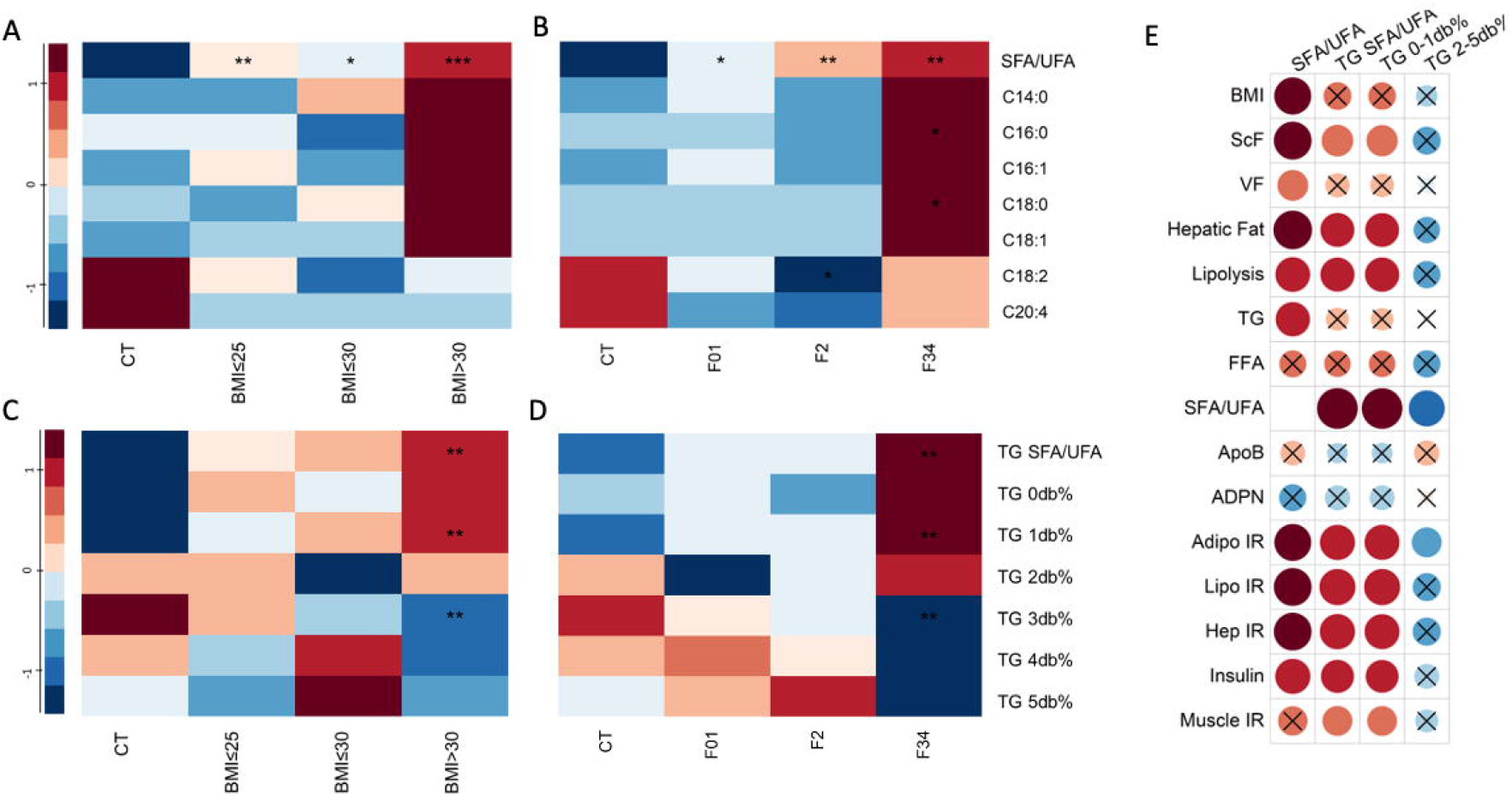
In panel A and B heatmaps of circulating FFA concentration levels and ratio of saturated to unsaturated FFA (SFA/UFA) in controls and NAFLD subjects divided by their BMI or degree of fibrosis, respectively. In panel C and D heatmaps of circulating TGs grouped in accordance with the numbers of double bonds and saturated to unsaturated TAG ratio in controls and NAFLD subjects divided by their BMI or degree of fibrosis, respectively. Values are reported as median within the groups, and normalization was applied to the rows. Specific p values for non-parametric Mann Whitney’s test vs control subjects: *<0.05, ** <0.01, ***<0.001, ****<0.0001. In panel E Pearson’s correlation matrix shows univariate association among the degree of saturation of FFA and TAG and metabolic parameters. Correlations that are not statistically significant (Pearson’s p-value ≥ 0.05) were crossed out.

We also observed that the saturated/unsaturated fatty acid ratio (in TG and FFA) was associated to increased Adipo-IR, Hep-IR, Muscle-IR and IHTG accumulation, and to a lesser extent to SC-AT but not to VF (**Fig. 4 E**), likely because the fatty acids released by VF into the portal vein are in great part taken up by the liver.

## Discussion

Abdominal obesity is a known risk factor for the development of NAFLD [33] since it’s strictly related to peripheral and Hep-IR [2]. The rate of FFA mobilization from the adipocytes embedded within the deep abdominal adipose tissue is higher compared to SC-AT [6, 42, 43], and previous studies suggested that VF is more strongly related to insulin resistance than SC-AT [6, 44, 45]. The impaired antilipolytic effect of insulin on adipose tissue and the FFA efflux from the VF directly to the liver through the preferential portal vein, affects liver function, thus promoting steatosis. But whether VF accumulation is the cause or the consequence of the onset and development of NAFLD is still being debated. In this study, we corroborate a strong correlation between VF accumulation and Hep-IR and IHTG accumulation, which occur also in lean NAFLD individuals without diabetes.

Although several studies demonstrated the involvement of VF in insulin resistance especially in the liver [24, 46-51], there are conflicting data in this regard. We have shown that fasting EGP and gluconeogenesis were associated only to increased VF while Hep-IR and impaired insulin mediated suppression EGP were associated to both VF and IHTG [24]. Fabbrini et al. questioned the role of VF, compared to IHTG, in the association with insulin resistance [52]; however, the group matched for IHTG with high/low VF had a mean hepatic fat of 13.3% [52], i.e. double the limit for NAFLD. A reanalysis of the data by grouping the subjects from low IHTG-Low VF to high IHTG-high VF showed that insulin resistance is increased proportionally with both visceral and liver fat [2]. The importance of high VF regardless of high or low IHTG was also confirmed in a recent study that showed how VF was associated to higher risk of type 2 diabetes [53]. The preferential accumulation of fat and calories in visceral rather than subcutaneous fat, can be related to adipose tissue expansion, either by a combination of an increase in adipocyte number (hyperplasia), or size (hypertrophy). While hyperplasia is associated with a beneficial and protective asset of adipose tissue, hypertrophy results in increased inflammation, macrophage infiltration, and reduced insulin sensitivity [54-56]. Whether VF is more prone to expand in a hypertrophic manner and develop inflammation compared to SC-AT is still unclear. It has been proposed that VF and IHTG accumulation are the consequence of a dysfunctional SC-AT that limits peripheral adipose tissue expansion. Treatment with pioglitazone ameliorates adipose tissue dysfunction and determines fat redistribution by decreasing both VF and IHTG and increasing SC-AT [57, 58]. Lipodystrophy is often study as a model of dysfunctional adipose tissue since it is associated with low subcutaneous fat and hepatic fat accumulation. Patients with generalized lipodystrophy have also low VF, while those with partial lipodystrophy have high VF [59]. Patients with severe lipodystrophy have also increased lipolysis and severe Hep-IR [60]. We have recently shown that the reduction of hepatic fat after pioglitazone treatment and the improvement in liver histology, was mediated by the decrease in VF and increase in SC-AT (ie by the VF/SC-AT ratio) [58]. These results indicate that the excess FFA that is not stored in the subcutaneous adipose tissue is then metabolized in the liver, thus increasing hepatic TG accumulation.

It has been reported that lean NAFLD individuals had a higher amount of daily calory intake compared to lean controls [61-63]. Notably, the Diabetes Remission Clinical Trial (DiRECT) study recently revealed a significant and sustained improvement of T2D remission and β-cell function after a very low-calorie diet in patients with early onset of the disease. Calorie restriction produced a substantial weight-loss, which was associated with a decrease in hepatic and intra-pancreatic fat content [64, 65]. Therefore, reducing calory intake may be an efficient strategy to reduce VF accumulation, thus avoiding the deleterious consequences that triggers the onset of Hep-IR and steatosis. The reason why an accumulation of VF occurs independent of weight-gain is not so clear. Some studies suggest that it may be related to stress associated with overeating or increased cortisol clearance. Genetics may also accentuate VF accumulation in response to stress [66]. Moreover, the characteristics of the subcutaneous adipose tissue and the expression of lipid storage-related genes determines the expansion of subcutaneous adipose tissue in response to excessive caloric intake [67].

Adipose tissue is not only a reservoir for energy storage and utilization, but also senses energy demands and secrete adipokines such as adiponectin, which exerts protective actions for liver injury by enhancing fatty acid β-oxidation, and reducing Hep-IR [68-71]. Studies on the association of adiponectin with VF rather than SC-AT have been conflicting [72-78]. Here, we showed that adiponectin levels were reduced in response to increased VF, and were inversely associated with increased Hep-IR, and steatosis. A significant decrease of circulating adiponectin was observed in lean NAFLD individuals compared to controls, thus suggesting that circulating levels of adiponectin as a potential biomarker to track development of NAFLD, VF accumulation and dysfunction. Thus, fat distribution is an important risk factor for the development and progression of NAFLD, because VF, unlike SC-AT, not only contributes to hepatic fatty acid overflow but it is significantly increased in the early stages of the disease.

Not only excessive caloric intake but also excess intake of saturated fat can be crucial in driving VF accumulation and liver damage [79] and also being associated to more severe forms of NAFLD [41]. It is already known that different classes of lipids, according to their degree of saturation, can impact metabolism. An increased proportion of unsaturated compared to saturated fatty acids, i.e., a decreased SFA/UFA ratio in animal models, has been associated, with a favorable serum lipid profile and activation of hepatic enzymes involved in antioxidative pathways [80]. A diet high in UFA and low in SFA decreases fat accumulation in white adipose tissue, and is associated with increased expression of hepatic lipolytic enzymes and enhanced fatty acid β-oxidation [81]. Target lipidomic analysis performed here revealed that novel lipid species may serve as early markers of liver damage such as SFA and saturated TG, independently of obesity. We also hypothesized that the SFA/UFA ratio could be considered as an index for the evaluation of metabolic health. Our results show that in NAFLD individuals the SFA/UFA ratio was significantly increased compared to healthy subjects indicating an imbalance in SFA in NAFLD, as previously suggested [12, 82-85]. These lipid alterations were strongly correlated with increased VF, Adipo-IR, Hep-IR and IHTG. These metabolic disturbances result in an increased demand for insulin secretion to ‘overcome’ insulin resistance, thus setting in place a vicious cycle. Indeed, insulin resistance is tethered to higher levels of insulin than expected relative to the level of glucose.

Although the results of this study are confirmatory, the main strength of our study is that we investigated non diabetic individuals with biopsy-proven NAFLD, i.e., excluding confounding factors strongly related to visceral adiposity, such diabetes and antidiabetic treatments, which influence fat accumulation and distribution [86-89]. Moreover, the individuals BMI ranged from 18-37 kg/m^2^ which allowed us to stratify individuals by BMI and explore the pivotal role of VF on IHTG accumulation and damage, independently of body weight. Moreover, the use of tracer studies provided a more thorough understanding of the association of VF with lipid and glucose fluxes, and IR in liver muscle and adipose tissue.

A limitation of the study is the low number of patients recruited to a single center, but this was due to difficulties and costs to perform tracer studies and MRI scans in a large number of subjects. In addition, we did not perform analysis stratified by sex, because the women recruited in this study were less than men. Therefore, we cannot ignore that sex differences can influence VF and SC-AT distribution, and correlation with HF content [42, 90]. Future studies on larger cohorts stratified for sex and BMI would be beneficial to study the onset and progression of lean NAFLD.

In conclusion, our findings indicate that high VF accumulation and saturated lipids as early indicators of severe NAFLD independently of BMI, therefore insinuating fat composition and distribution as key players in the regulation of lipid metabolism and NAFLD progression. A deeper investigation of fat distribution would be instrumental in defining the best targeted-interventional approach and to characterize patients at increased risk of disease development.

## Abbreviations

NAFLD: Nonalcoholic fatty liver disease
AT: Adipose tissue
IHTG: Intrahepatic triglyceride
IR: Insulin resistance
SC-AT: Subcutaneous fat
VF: Visceral fat
FFAs: Free fatty acids
TAG: Triglyceride
MS: Mass spectrometry
NASH: Non-alcoholic steatohepatitis
MRI: Magnetic resonance imaging
HDL: High density lipoprotein
LDL: Low density lipoprotein
LFTs: Liver function tests
APO-B: Apolipoprotein B
APO-A1: Apolipoprotein A1
NAS: NAFLD activity score
ROI: Region of interest
LFF: Liver fat fraction
EGP: Endogenous glucose production
Hep-IR: Hepatic insulin-resistance
Muscle-IR: Muscle insulin-resistance
Adipo-IR and Lipo-IR: Adipose tissue insulin-resistance
ALT: Alanine aminotransferase
AST: Aspartate aminotransferase
GGT: Gamma-glutamyltransferase

## Acknowledgments and Funding

A.G. and E. B. received financial support from the European Union’s Horizon 2020 for the project EpoS: Elucidating Pathways of Steatohepatitis (Grant agreement ID: 634413). A.G. received financial support from the European Union’s Horizon 2020 Research and Innovation Programme under the Marie Skłodowska-Curie Grant Agreement ID 722619 (FOIE GRAS project) and Grant Agreement ID 734719 (mtFOIE GRAS project)

---

**Graphical abstract.**
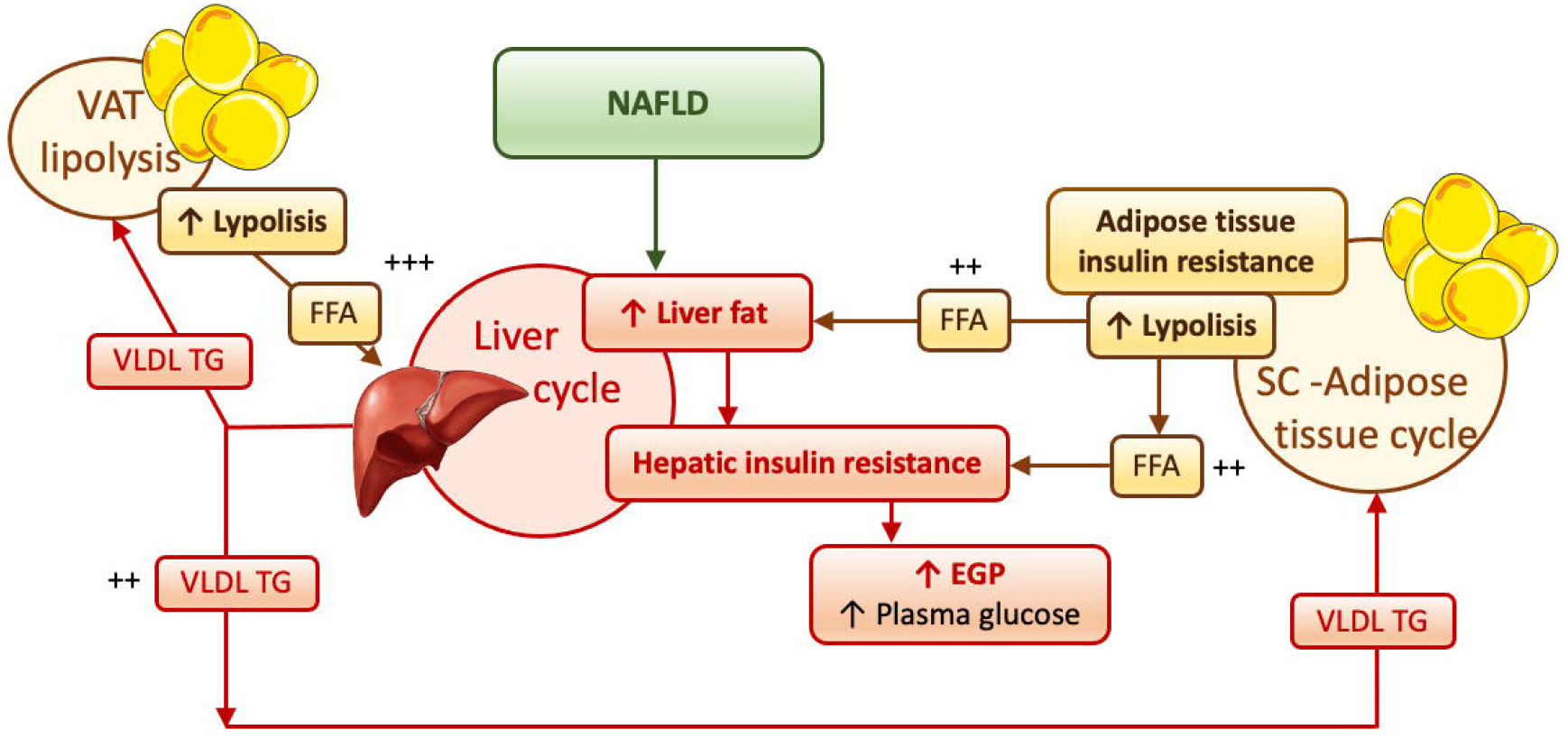
NAFLD and its progression are associated with altered fat distribution, and adipose tissue and hepatic insulin resistance. NAFLD is an heterogenous metabolic disorder that affects several organs including the liver, adipose tissue, and the way these organs interact with each other. Excess of fat accumulation, both visceral and subcutaneous, and the onset of adipose tissue insulin resistance, with consequent increase in lipolysis and efflux of fatty acids to the liver, leads to ectopic accumulation and the establishment of hepatic insulin resistance. In turn, the increased synthesis and secretion of VLDL from the liver results in an overload to the adipose tissue, exacerbating its dysfunction.

